# A sparse negative binomial classifier with covariate adjustment for RNA-seq data

**DOI:** 10.1101/636340

**Authors:** Tanbin Rahman, Hsin-En Huang, An-Shun Tai, Wen-Ping Hsieh, George Tseng

**Affiliations:** University of Pittsburgh; National Tsing Hua University

## Abstract

Supervised machine learning methods have been increasingly used in biomedical research and in clinical practice. In transcriptomic applications, RNA-seq data have become dominating and have gradually replaced traditional microarray due to its reduced background noise and increased digital precision. Most existing machine learning methods are, however, designed for continuous intensities of microarray and are not suitable for RNA-seq count data. In this paper, we develop a negative binomial model via generalized linear model framework with double regularization for gene and covariate sparsity to accommodate three key elements: adequate modeling of count data with overdispersion, gene selection and adjustment for covariate effect. The proposed method is evaluated in simulations and two real applications using cervical tumor miRNA-seq data and schizophrenia post-mortem brain tissue RNA-seq data to demonstrate its superior performance in prediction accuracy and feature selection.

## 1 Introduction

In the past two decades, microarray and RNA sequencing (RNA-seq) are routine procedures to study transcriptome of organisms in modern biomedical studies. In recent years, RNA-seq [5, 20] has become a popular experimental approach for generating a comprehensive catalog of protein-coding genes and non-coding RNAs [13], and it largely replaces the microarray technology due to its low background noise and increased precision. The most important difference between RNA-seq and microarray technology is that RNA-seq outputs millions of sequencing reads rather than the continuous fluorescent intensities in microarray data. Unlike microarray, RNA-seq can detect novel transcripts, gene fusions, single nucleotide variants, and indels (insertion/deletion). It can also detect a higher percentage of differentially expressed genes than microarray, especially for genes with low expression [24].

In machine learning, classification methods are used to construct a prediction model based on a training dataset with known class label so future independent samples can be classified with high accuracy. For example, labels in clinical research can be case/control, disease subtypes, drug response or prognostic outcome. Many popular machine learning methods have been widely applied to microarray studies, such as linear discriminant analysis [9], support vector machines [3] and random forest [7]. However, for discrete data nature in RNA-seq, many powerful tools for microarray assuming continuous data input or Gaussian assumption may be inappropriate. A common practice to solve this problem is to transform RNA-seq data into continuous values such as FPKM or TPM [6] and possibly taking additional log-transformation. However, such data transformation can lead to loss of information from the original data [14, 18], producing less accurate inference. Particularly, the transformation often produces greater loss of information for genes with lower counts [15]. To accommodate discrete data in RNA-Seq, Poisson distribution and negative binomial distribution are two common distributions expected to better fit the data generation process and data characteristics. Witten [22] proposed a sparse Poisson linear discriminant analysis (sPLDA) based on Poisson assumption for the count data. However, Poisson distribution assumes equal mean and variance, which is often not true. In real RNA-seq data, the variance is often larger than the mean, leading to the need of an overdispersion parameter. Witten [22] reconciled this problem by proposing a power transformation to the data for eliminating overdispersion. However, as we will see later, the power transformation can perform well when the overdispersion is small but performs poorly when overdispersion becomes large. Hence, direct modeling by negative binomial assumption rather than a Poisson distribution is more appropriate. To this end, Dong et al. [8] proposed negative binomial linear discriminant analysis (denoted as NBLDA_PE_) by adding a dispersion parameter. They, however, borrowed the point estimation from sPLDA in [22] and did not pursue a principled inference such as maximum likelihood, consequently producing worse performance than the method we will propose later.

Since the number of genes is often much larger than the number of samples in transcriptomic studies (a standard “small-n-large-p” problem), feature selection is critical to achieve better prediction accuracy and model interpretation. Witten [22] proposed a somewhat ad hoc soft-thresholding operator, similar to univariate lasso estimator in regression, for gene selection in sPLDA but the method is not applicable to the NBLDA_PE_ model due to the addition of dispersion parameter. In the NBLDA_PE_ model proposed by [8], feature selection issue was not discussed, except that they used “edgeR” package to reduce the number of genes in the input data. Such a two-step filtering method is well-known to have inferior performance than methods with embedded feature selection. In fact, Zararsiz et al. [23] have compared sPLDA and NBLDA_PE_, and showed that the power transformed sPLDA generally performed better than NBLDA_PE_ in their simulations and the worse performance in NBLDA_PE_ mainly came from the lack of feature selection. Finally, another critical factor to consider in transcriptomic modeling is the adjustment of covariates such as gender, race and age since it is well-known that many genes are systematically impacted by these factors. For example, Peters et al. [16] have identified 1,497 genes that are differentially expressed with age in a whole-blood gene expression meta-analysis of 14,983 individuals. A classification model allowing for covariate adjustment is expected to provide better accuracy and deeper biological insight.

To account for all aforementioned factors, we propose a sparse negative binomial model for classification analysis with covariate selection and adjustment. The method is based on generalized linear model (GLM) with a first regularization for feature sparsity. The GLM framework also allows straightforward covariate adjustment and a second regularization term on covariates, facilitating further covariate selection. Such covariate adjustment is not possible through existing sPLDA or NBLDA_PE_ methods. The paper is structured as following. In Section 2.1, we will briefly describe the two existing methods sPLDA [22] and NBLDA_PE_ [8], and then followed by our proposed methods sNBLDA_GLM_ and sNBLDA_GLM.sC_ in Section 2.2. Section 2.3 and 2.4 will discuss parameter estimation and model selection of the proposed method. Benchmarks for evaluation are described in Section 2.5. Section 3 presents simulation studies and Section 4 shows two real applications of cervical tumor miRNA data and schizophrenia RNA-seq data. Conclusion and discussion are included in Section 5.

## 2 Existing and proposed methods

In this section, we will first describe two existing methods for classification analysis of count data from RNA-seq and then propose our new method. To unify the notation, denote by **X** the count data matrix with elements *X*_*ij*_ referred to the sequence count for the *j*-th gene and the *i*-th sample (*i* = 1, 2, … *n* and *j* = 1, 2, … *p*). In addition, **x_i_** = (*X*_*i*1_ … *X*_*ip*_)^*T*^ denotes *i−*th row of **X**, corresponding to feature measurements of observation *i*. Also, define 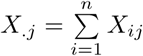, 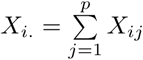 and *X_.._* = ∑_*i,j*_ *X*_*ij*_. Moreover, in the classification setting where each observation belongs to one of the *K* classes, we let disjoint sets *C*_*k*_ ⊂ {1, …, *n*} contain the indices of observations in class *k*. That is, class label *y*_*i*_ = *k* if and only if *i* ∈ *C*_*k*_. Furthermore, we denote Furthermore, we denote 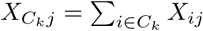.

### 2.1 Two existing methods for classification of RNA-seq data

#### 2.1.1 Sparse Poisson linear discrimination analysis (sPLDA)

Witten [22] introduced a log-linear Poisson model with feature selection, which resulted in a simple diagonal linear discriminant analysis suitable for count data (referred as “sPLDA” hereafter in this paper). Under the assumption of gene independence, the model is based on the following formulation,

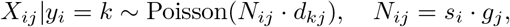

where *s*_*i*_ is the normalizing factor (a.k.a. size factor) for sample *i* and *g*_*j*_ is the ground mean for the *j*-th gene, allowing for variations both in samples and genes. For a given gene *j*, *d*_1*j*_, …, *d*_*kj*_ allows the *j*-th gene to be differentially expressed between the classes if any of *d*_*kj*_ ≠ 1(1 ≤ *k ≤ K*).

RNA-Seq data often contain over-dispersion such that variances are larger than means, whereas an important constraint in Poisson model is the equivalent mean and variance. To overcome this, Witten [22] proposed a transformation of count data 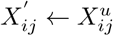 with a proper choice of *u* such that 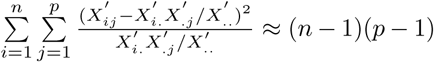. From simulations of the original paper, this correction performs well in the presence of weak to moderate overdispersion.

Suppose 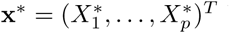 be a future new sample for prediction. The discriminant score for assigning **x**\* to class *k* is,

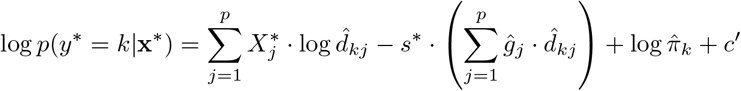

where *y*\* is the predicted label, 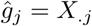, 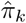 is the estimate of prior probability of belonging to the *k*th class estimated by the fraction of samples belonging to class *k* and *s*\* is the estimated normalization factor for the new sample **x**\* for which we do not know the class label. The classifier assigns **x**\* to the class with the largest discriminant score. The paper also implemented a somewhat ad hoc soft-thresholding operator for feature selection in the classifier, which is motivated from univariate lasso regularization in regression for feature selection: 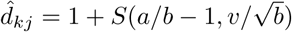, where *a* = *X*_*Ckj*_+ *β*, 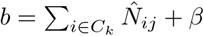, *β* is the hyperparameter pre-determined in the estimation of *d*_*kj*_, *v* is the tuning parameter chosen by cross validation and *S*(*x, a*) = sign(*x*)(*|x| − a*)_+_ is the soft thresholding parameter. 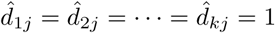 means gene *j* is not differentially expressed across the classes and thus, is not selected in the classifier.

#### 2.1.2 Negative binomial linear discrimination analysis (NBLDA_PE_)

Dong et al. [8] extended sPLDA into a negative binomial model to explicitly allow overdispersion property in RNA-seq data:

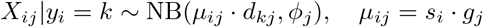

Under the formulation, *E*(*X*_*ij*_) = *µ_ij_* and 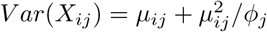. Similar to sPLDA, for a new observation **x**\*, prediction is made by the maximized discriminant score:

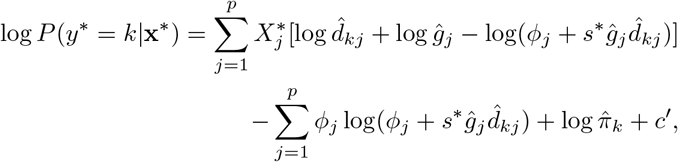

where *ϕ_j_* is the dispersion parameter for the jth gene, 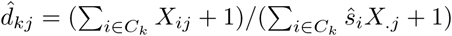 and 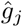 is the same as defined previously. We note that the point estimate of 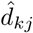 and 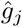 are borrowed directly from Witten’s sPLDA model without theoretical justification and the similar soft-thresholding in sPLDA cannot be easily incorporated into the procedure due to the increased complexity with *ϕ_j_*.

In the literature, several popular procedures have been used for estimating the size factor, including simple sum of counts, median ratio [1] and quantile method [4]. Witten [22] and Dong et al. [8] showed that the performance is comparable among the three methods. Here, we will use the quantile method for all methods for a fair comparison. In quantile method, the normalization factor for sample *i* (1 ≤ *i* ≤ *n*) is estimated as 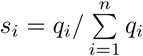 (or equivalently some papers also use 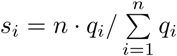, which is what we adopt in this paper), where *q*_*i*_ is the 75th quantile of sequence counts of all genes for the *i*th sample. For a new sample **x**\*, the normalizing factor is estimated as 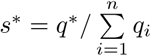, where *q*_*i*_ (1 ≤ *i* ≤ *n*) come from training data and *q*\* is the 75th count quantile for sample **x**\*. Note that the vector of normalization factors and dispersion denoted by **s** and ***ϕ*** respectively will be pre-estimated in all negative binomial models in this paper before inference. ***ϕ*** are estimated by weighted likelihood empirical Bayes method using the edgeR package [17] with class label considered. We denote the method proposed by [8] as “NBLDA_PE_” to emphasize the ad hoc “point estimation” procedure inherited from sPLDA in [22].

### 2.2 Proposed method: sparse negative binomial classifier via generalized linear model

We first consider a model without covariate in section 2.2.1. Then we extend it to covariate in section 2.2.2.

#### 2.2.1 Sparse negative binomial classifier without covariate adjustment (sNBLDA_GLM_)

Similar to NBLDA_PE_, we specify the following negative binomial model in a generalized linear model (GLM) setting:

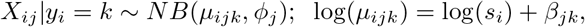

where *s*_*i*_ is the normalization factor of the *i*-th sample, *β_jk_* is the mean count in log-scale of the *k*-th class for the *j*-th gene and *ϕ_j_* is the dispersion parameter of the *j*-th gene. Under the assumption of independence between genes, the corresponding log-likelihood can be written as,

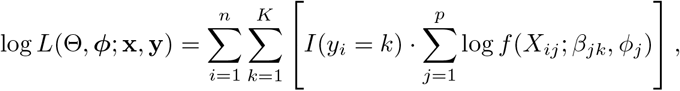

where, Θ = {(***β***_*k*_, ***ϕ***); *k* = 1, …, *K*}, ***β***_*k*_ = (*β*_1*k*_, …, *β*_*pk*_), ***ϕ*** = (*ϕ*_1_, …, *ϕ*_*p*_), *I*(*y*_*i*_ = *k*) is the indicator function taking value 1 if *y*_*i*_ = *k* and 0 otherwise, and *f* (*X*_*ij*_; *β*_*jk*_, *ϕ*_*j*_) is the density function of NB(*s*_*i*_ exp (*β*_*jk*_), *ϕ*_*j*_). Now, suppose we have a new observation **x**\* for which we intend to predict the class label. By Bayes theorem, we can derive the discriminant score as

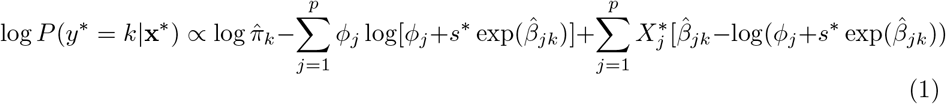

Here, **x**\* is assigned to class *k* for which the discriminant score is maximized. Note that the form of the discriminant score in the current model is identical to that proposed in [8], except that we reparametrize *µ*_*ijk*_ = *s*_*i*_*g*_*j*_*d*_*kj*_ to log(*µ*_*ijk*_) = log(*s*_*i*_) + *β*_*jk*_. The major difference is in the parameter estimation. [8] directly borrows the point estimation of *µ*_*ijk*_ from the Poisson model in [22], while we will derive MLE of Equation (2) (see below) using iteratively reweighted least squares (IRLS) method to be shown in the next subsection.

In order to incorporate variable (gene) selection, we add a penalty term 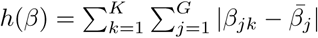. Here, 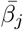 is the average of *β*_*jk*_’s over the *K* classes for a given *j*-th gene. Hence, the following penalized likelihood is maximized to obtain estimation of *β* with pre-estimated ***ϕ***:

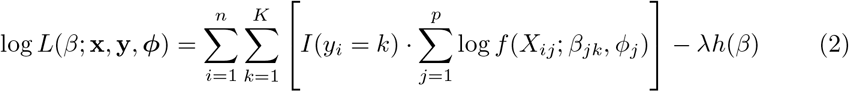

Here, *β* is the collection of all *β*_*jk*_ parameters and *λ* is a tuning parameter controlling sparsity of the variable selection. The form of the discriminant scores for prediction is the same as in equation 1.

#### 2.2.2 Sparse negative binomial classifier with covariate adjustment (sNBLDA_GLM.C_ and sNBLDA_GLM.sC_)

In real applications, information of multiple clinical variables is often available and some of them may be associated with subsets of genes. Commonly encountered clinical variables can include age, gender, race, etc. Failure of covariate adjustment can greatly reduce prediction accuracy and replicability. In our GLM framework, covariate adjustment can be straightforwardly incorporated in the linear regression term:

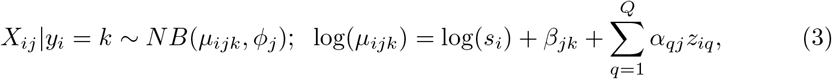

Here, **z**_**q**_ = (*Z*_1*q*_, …, *Z*_*nq*_) includes values of the *q*-th covariate over *n* samples and parameter *α*_*qj*_ corresponds to the coefficient of the *q*-th covariate in the *j*-th gene. Under the assumption of gene independence and adding penalty terms for both genes and covariates, the problem can be presented as maximization of the following penalized log-likelihood with double regularization:

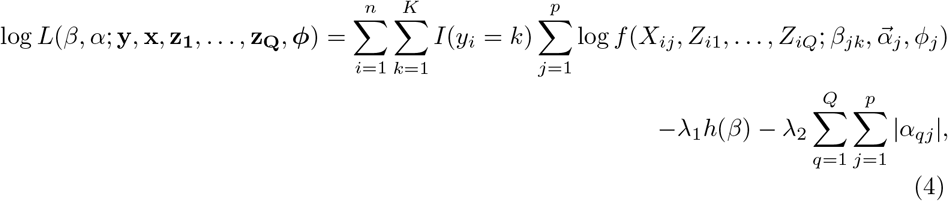

where, *β* is the collection of all *β*_*jk*_ parameters and *α* is the collection of all *α*_*qj*_ parameters. *λ*_1_ and *λ*_2_ are tuning parameters controlling for levels of sparsity of variable selection in genes and covariates, respectively.

Similarly, for a new sample **x**\* with vector of clinical vector 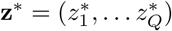 under this framework, we can derive the following discriminant score:

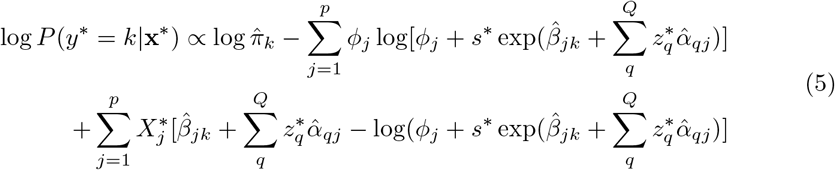

As before, **x**\* is assigned to the class with the highest discriminant score. We note that when *λ*_2_ = 0, Equation 4 performs covariate adjustment using all covariates for all genes without regularization in covariate parameters *α*_*qj*_. We will denote this method as “sNBLDA_GLM.C_”. In this case, when the number of covariates *Q* becomes large, performance of parameter estimation and prediction accuracy are expected to decline. With proper choice of *λ*_2_ in Equation (4), the method can adequately select a subset of covariates in each gene to improve the performance. For illustration purpose, we refer to this method as “sNBLDA_GLM.sC_” in this paper, where “sC” means sparsity on covariates.

### 2.3 Estimation in sNBLDA_GLM_ and sNBLDA_*GLM.sC*_

#### 2.3.1 Estimation of sNBLDA_GLM_

Maximizing the log-likelihood derived in Equation (2) is equivalent to minimizing the following penalized weighted least square function,

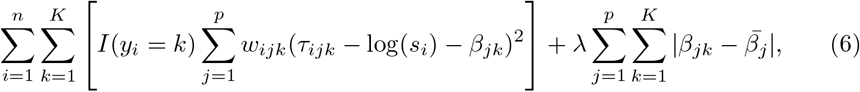

where 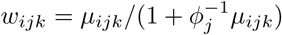 and *τ*_*ijk*_ = log(*s*_*i*_) + *β*_*jk*_ + (*x*_*ij*_ − *µ*_*ijk*_)/*µ*_*ijk*_. Given the estimates at the t-th step, the updates of (t+1) step is:,

1. Calculate 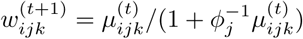
2. Update 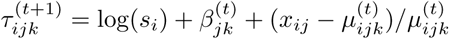
3. Solve 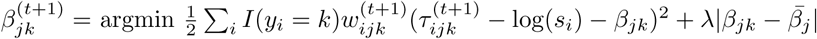
4. Update 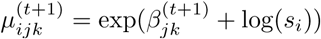

This is repeated until convergence of 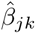. The update of 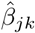 in Step (3) is given by,

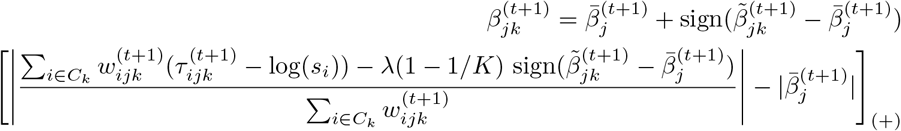

Here, [ ]_(+)_ is soft thresholding function such that [*u*]_(+)_ takes the value *u* when *u* is positive and 0 otherwise, 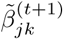 is the estimate of *β*_*jk*_ under no penalization and 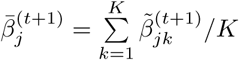.

#### 2.3.2 Estimation of *sNBLDA_GLM.sC_*

Similar to sNBLDA_GLM_, the problem of maximizing the penalized log-likelihood in Equation (4) can be represented as minimizing the penalized weighted least square function given below in Equation (7),

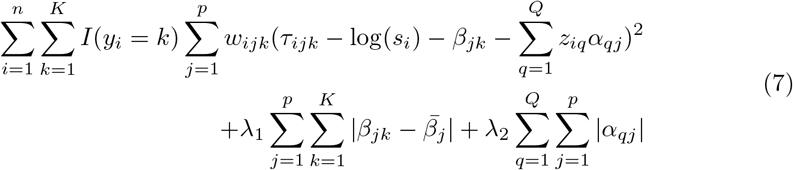

where, 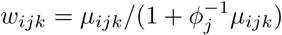 and 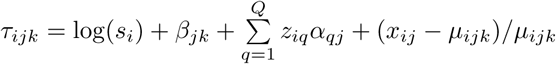. The estimation of each of the 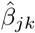 and *α*_*qj*_ is given by the following algorithm. The steps involved in IRLS given the estimates obtained at the *t*-th step is given below,

1. Calculate 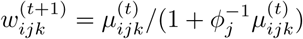
2. Update 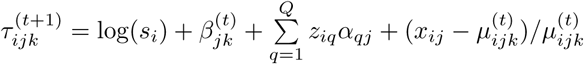
3. Solve 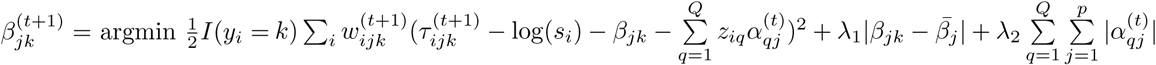
4. Solve 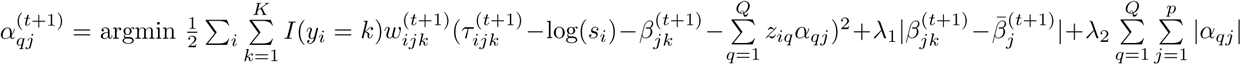
5. Update 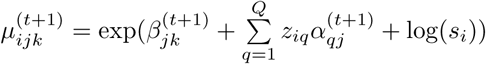

The steps are repeated until convergence of the parameters *β*_*jk*_, *α*_1*j*_, …, *α*_*qj*_. Then the penalized estimate of the parameters in step 3 and step 4 are respectively given by,

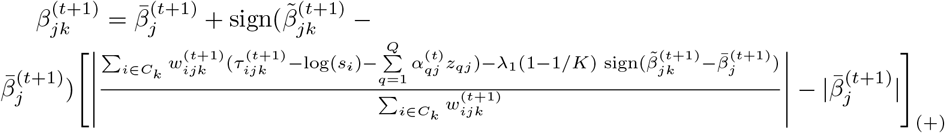

and, 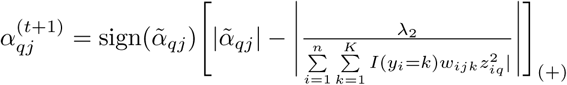 where,

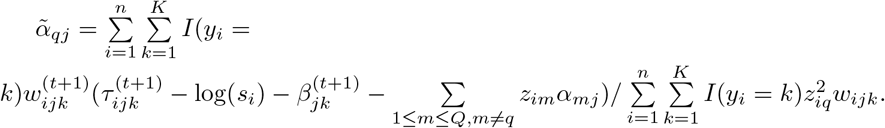

### 2.4 Selection of tuning parameters in regularization

Both sNBLDA_GLM_ and sNBLDA_GLM.sC_ methods involve selection of regularization parameters *λ* or (*λ*_1_*, λ*_2_). We apply V-fold cross validation as a tool to determine the tuning parameter [19]. For each given tuning parameter, we divide the dataset into *V* equal folds and samples in the *K* classes are split into *V* folds as even as possible. In each iteration, one fold is set aside as the test set and the remaining (*V* − 1) folds are used as the training set. The classifier is built from the training set and then validated in the test set for evaluating accuracy. This procedure is repeated until all *V* folds have been chosen as the test set and the averaged accuracy is calculated. The tuning parameter corresponding to the highest averaged accuracy is chosen for the final model construction. We apply 10-fold (V=10) cross validation for all simulations and real applications in this paper. We note that nested cross validation is used for real applications for a fair accuracy evaluation. In this case, the outer loop of 10-fold cross validation is conventionally used to estimate accuracy. In each cross validation, the 9 folds of training set undergo an inner loop of 10-fold cross validation to determine *λ* or (*λ*_1_*, λ*_2_).

### 2.5 Benchmarks for evaluation

Performance of different methods will be judged by two major criteria: accuracy of prediction and accuracy of feature selection. For prediction performance, simple averaged accuracy is used when true class labels are known: 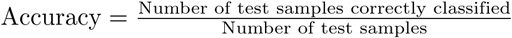. For feature selection performance, we derive the area under the curve (AUC) [2] values of the receiver operating characteristic (ROC) curves. We also evaluate the performance of sNBLDA_GLM_, sNBLDA_GLM.sC_ and sPLDA in terms of estimating the true parameters *β*_*jk*_ when the gene expression is affected by covariates. Here, we define 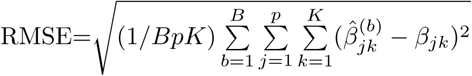 where *B* is the number of datasets simulated.

## 3 Simulations

In this section, we will devise two simulation schemes to compare the performance of sPLDA and NBLDA_PE_ to our proposed model sNBLDA_GLM_ and sNBLDA_GLM.sC_ under different settings. In Simulation 1, there is no covariate effect over the expression levels of the genes. Here, we compare sPLDA, NBLDA_PE_ and sNBLDA_GLM_ over different level of signal strength under three different levels of dispersion in the data. In Simulation 2, we develop a simulation scheme where two covariates are introduced which can affect expression level of certain proportion of the genes. Here, we compare sPLDA, NBLDA_PE_, sNBLDA_GLM_ and sNBLDA_GLM.sC_ in the presence of covariate effects.

In order to mimic real data, we use a real RNA-seq dataset downloaded from Gene Expression Omnibus (GEO, GSE47474) to retrieve key parameters and perform the simulation. The dataset includes 72 samples with 36 coming from HIV-1 transgenic and 36 from control rat strains [12]. We compute the mean counts of each gene over all samples to obtain an empirical distribution of mean counts, which will be used for obtaining baseline expression levels in all the simulations. Each simulation is repeated 100 times and the average result is reported.

### 3.1 Simulation settings

#### Simulation 1: Without covariate effect

In this simulation, we sample the count data by *X*_*ij*_|*C*_*i*_ = *k* ~ *NB*(*s*_*i*_*b*_*j*_ exp (*δ*_*jk*_∆_*j*_), *ϕ*_*j*_) for each gene *j*(1 ≤ *j* ≤ 1000) and sample *i*(1 ≤ *i* ≤ 120) in class *k*(1 ≤ *k* ≤ 3), where the number of informative feature is 300. The notation of the parameters as well as the settings are given below:

- The library size factor *s*_*i*_ is sampled from Unif(0.75,1.25) for each sample *i*.
- *b*_*j*_ is the baseline which is sampled from the empirical distribution of the mean expression described previously.
- *δ*_*jk*_ represents the pattern of genes *j* in class *k*. For all *δ*_*jk*_ ∈ {−1, 0, 1}, 1 indicating a up-regulated trend of genes in this class relative to other classes, −1 indicating it is down-regulated and 0 indicating no difference.
- There exists three gene patterns for the 300 informative genes: (*δ*_*j*1_, *δ*_*j*2_, *δ*_*j*3_) = (1,0,−1), (0,1,1) and (−1,−1,0). For non-informative genes, the pattern is (0,0,0).
- Sample the main effect size parameter ∆_*j*_ for each gene *j* from a truncated normal distribution *TN* (*ζ*, 0.1^2^*, ζ/*2,∞), where *ζ* is the mean and values smaller than *ζ/*2 are truncated.
- *ϕ*_*j*_ ∼ *TN* (*ν*, 0.1, 0, ∞) and *ν* is chosen as 1, 5 and 10.
- 100 of the samples are used as training set and the remaining 1,000 samples are used as testing set

#### Simulation 2: Incorporating covariate effect

We sample the count data by 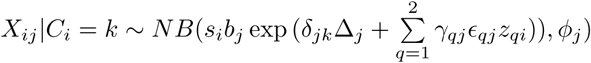 for each gene *j*(1 *≤ j ≤* 1000) and sample *i*(1 *≤ i ≤* 120) in class *k*(1 *≤ k ≤* 3) with two covariates (*z*_1_ and *z*_2_; Q=2), where the number of informative feature is 300. The notation of parameters are as follows:

- We generate a binary covariate (e.g. gender) for each sample *i* from *Ber*(0.5)(*i.e.z*_1*i*_ ~ Ber(0.5)) and generate a continuous covariate (e.g. age) for each sample *i* from *Gamma*(5, 10)
- *ϕ*_*j*_ ∼ *TN* (*ν*, 0.1, 0, ∞) where *ν* ∈ {10, 1}
- γ_*qj*_ represents the pattern of gene *j* in covariate *q* for all *γ*_*qj*_ ∈ {0, 1}; there exist three patterns: (*γ*_1*j*_, *γ*_2*j*_) = (1, 1), (1, 0), (0, 1), and (0, 0) with probablity (*ρ*/3,*ρ*/3,*ρ*/3 and 1-*ρ*) respectively. When *ρ* = 0, all genes are not impacted by covariates. We choose the proportion of covariate-impacted genes *ρ* to be 0.125, 0.25 and 0.5.
- Sample the main effect size parameter ∆_*j*_ for each gene *j* in class *k* from a truncated normal distribution *TN* (0.25, 0.1^2^, 0.125, ∞)
- The effect size parameter of covariates *ε*_*qj*_ for each gene *j* in covariate *q* is drawn from a truncated normal distribution *TN* (*η*, 0.1^2^*, η/*2,∞) × *κ* where *κ* takes value 1 with probability 0.5 and −1 otherwise. We use the different value of *η* ∈ {0.1, 0.3, 0.5, 0.7} for different level of signal strength of the covariates.
- Other parameters are set the same as Simulation 1 except that *ζ* is set at 0.25.
- 100 of the samples are used as training set and the remaining 1,000 samples are used as testing set.

### 3.2 Simulation results

Results of Simulation 1 are summarized in Figure 1. In Figure 1(a), average prediction accuracy of the three models sPLDA, NBLDA_PE_ and sNBLDA_GLM_ were compared over three different levels of dispersions *ν* ∈ {1, 5, 10}. The larger the value of *ν*, the smaller the level of dispersion in the simulated datasets. In all different levels of *ζ* and *ν*, sNBLDA_GLM_ outperformed the other two methods. As expected, NBLDA_PE_ was superior to sPLDA when *ν* was small (large overdispersion) but their performances became comparable when *ν* was large, confirming good performance of power transformation to correct dispersion in sPLDA only for small overdispersion. Figure 1(b) shows results of variable selection by AUC. sNBLDA_GLM_ clearly outperformed sPLDA in all cases while NBLDA_PE_ could not perform variable selection and was not applicable in this plot. The new method was also compared to three popular classification methods such as support vector machines (SVM), random forest (RF) and classification and regression tree (CART) in supplement Figure S4. The result showed inferior performance in these methods due to ignorance of count data and transformation to continuous inputs.

**Figure 1.**
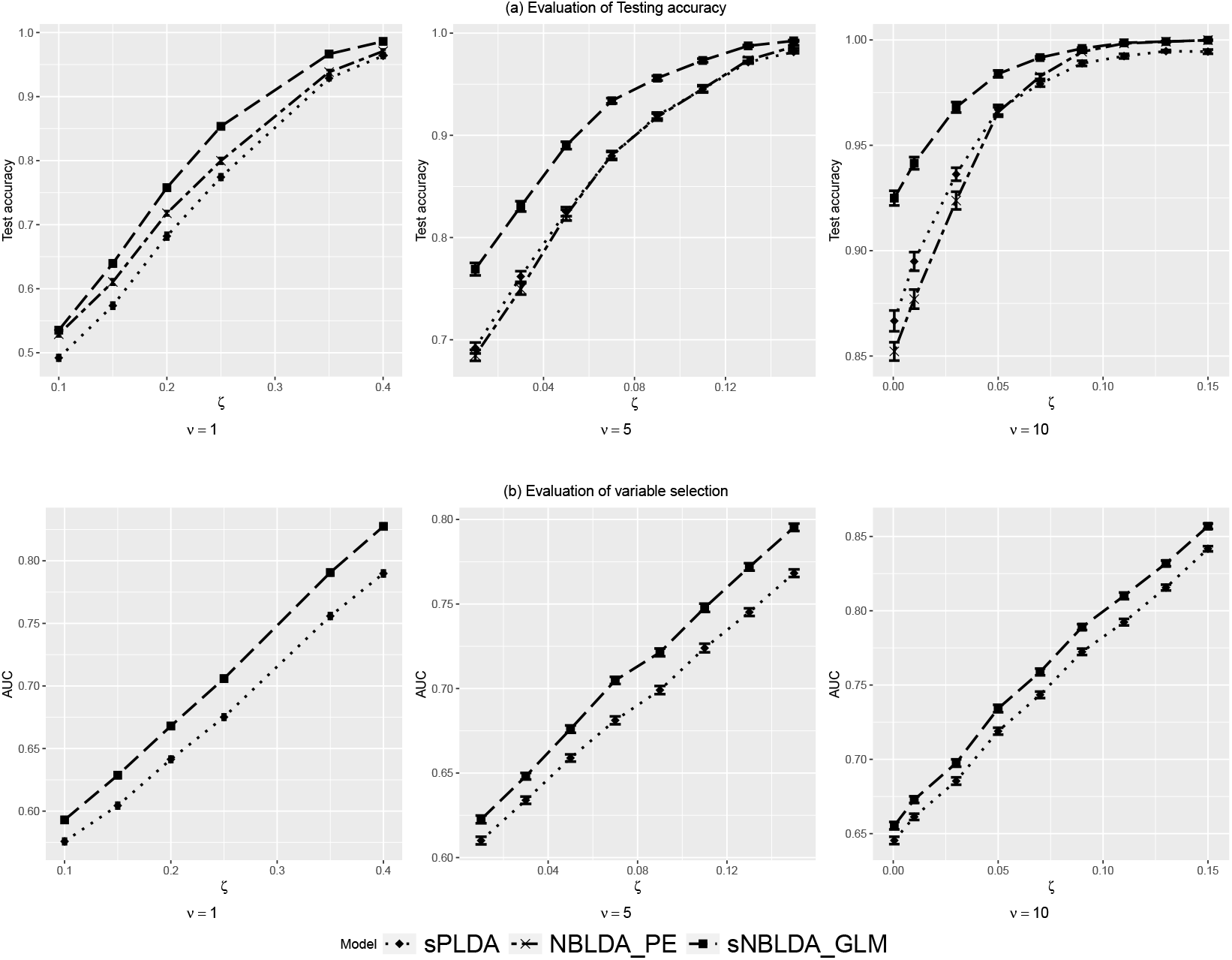
Results for Simulation 1 without covariate effect

Figure 2 demonstrates results of Simulation 2 using sPLDA, NBLDA_PE_, sNBLDA_GLM_ (no covariate adjustment) and sNBLDA_GLM.sC_ (with covariate adjustment and regularization) when varying percent of genes impacted by covariates *ρ* = 0.125, 0.25 and 0.5. Figure 2(a) shows averaged prediction accuracy of varying *η* and *ν* = 1 or 10. When *ν* = 1 (high level of dispersion), sNBLDA_GLM.sC_ outperformed all other methods as the impact of covariates on gene expression *η* increased. The prediction accuracy for sNBLDA_GLM.sC_ remained high with increased *η* due to its capacity of adjusting covariate effect, while prediction accuracy of the other three methods dropped with increased *η* although sNBLDA_GLM_ still outperformed sPLDA and NBLDA_PE_. When *ν* = 10, similar pattern was observed. The margin between sNBLDA_GLM.sC_ and sNBLDA_GLM_ became much smaller but sNBLDA_GLM.sC_ was still the best performer. Supplementary Figure S5 includes comparison with SVM, RF and CART, all of which performed much worse than sNBLDA_GLM.sC_.

**Figure 2.**
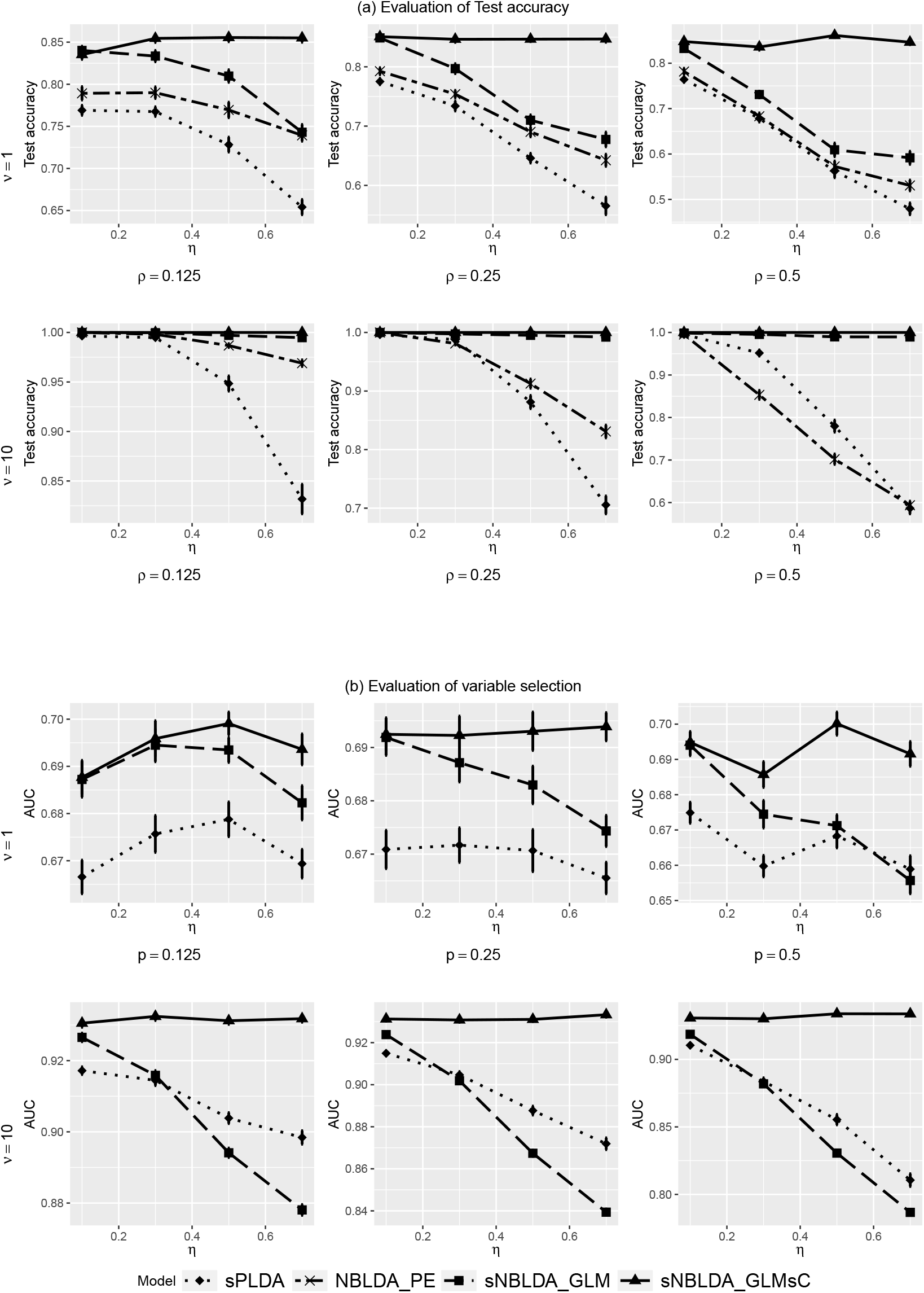
Results for Simulation 2 with covariate effect

Variable selection performance between sPLDA, sNBLDA_GLM_ and sNBLDA_GLM.sC_ is shown in 2(b). Similarly, we observed stable and high performance of sNBLDA_GLM.sC_ with increasing *η*, while performance of sNBLDA_GLM_ dropped for increased *η* due to the lack of covariate adjustment. sPLDA performed the worst in all cases under high level of dispersion. However, when the covariate effect is strong under moderate dispersion, it is seen to outperform NBLDA_GLM_ in terms of variable selection. It is intriguing that the variable selection gap between NBLDA_GLM.sC_ and NBLDA_GLM_ was larger in *ν* = 10 than in *ν* = 1, which is contrary to the prediction accuracy in Figure 2(a). An evaluation of the parameter estimates between sPLDA, sNBLDA_GLM_ and sNBLDA_GLM.sC_ was carried out in terms of RMSE in supplement Figure S1, where sNBLDA_GLM.sC_ performed the best. To examine the advantage of covariate regularization, we compared sNBLDA_GLM.C_ (i.e. *λ*_2_ = 0 in Equation 4; all covariates are used) with sNBLDA_GLM.sC_ (with covariate regularization) in Supplement Figure S2. The result shows clear improvement of covariate regularization on prediction accuracy but less on feature selection.

## 4 Real applications

### 4.1 Cervical tumor miRNA data

This RNA-seq dataset measured expression level of miRNAs in tumor and nontumor cervical tissues in human samples [21]. The dataset contains information of over 714 microRNAs for 29 control samples (samples with no tumor) and 29 tumor samples. No clinical information (covariates) is available for adjustment. This dataset has been used in both sPLDA and NBLDA_PE_ papers and thus is a good dataset to evaluate our new method. Dong et al. [8] found that NBLDA_PE_ performed better than sPLDA in terms of prediction accuracy because of high dispersion estimate in this dataset. In Figure 3,we compare prediction accuracy (y-axis) between sPLDA and sNBLDA_GLM_ based on 10-fold cross-validation when different number of genes are selected (x-axis) as proposed for the corresponding models. Since there is no variable selection in NBLDA_PE_, we only performed cross-validation considering all miRNAs (shown as “X” in the figure). sNBLDA_GLM_ generally outperformed the other two methods in different number of selected genes. Specifically, it achieved 95% prediction accuracy with a small number of 37 genes while NBLDA_PE_ and sPLDA achieved around 91% accuracy. Although the improvement in accuracy is marginal given the small sample size, the result indicates a trend of improvement of sNBLDA_GLM_ in prediction accuracy and variable selection.

**Figure 3.**
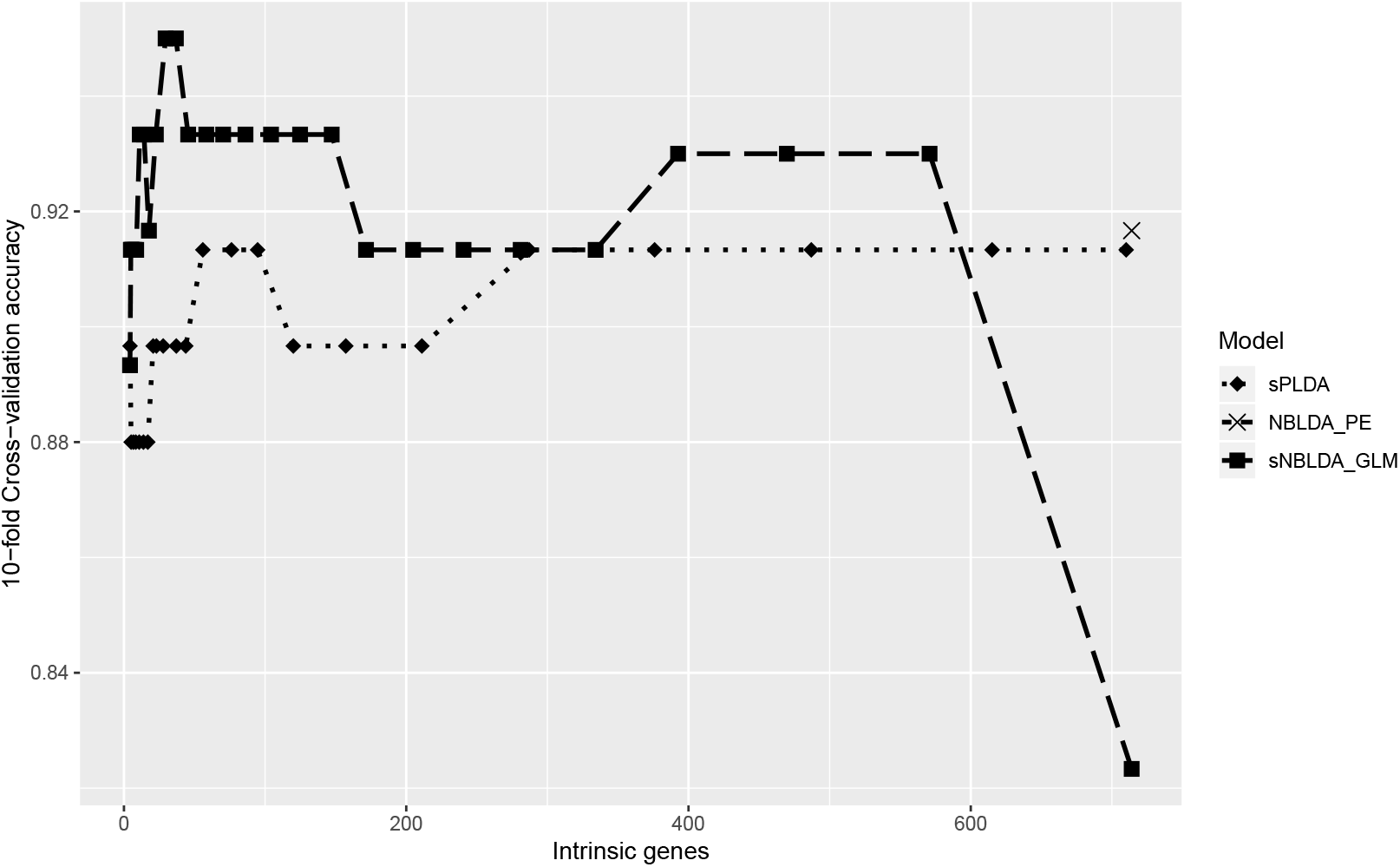
Prediction accuracy (y-axis) of sPLDA (dotted line) and sNBLDA_GLM_ (dashed line) with varying number of selected miRNAs (x-axis) in the cervical tumor application. NBLDA_PE_ does not allow variable selection and is shown with “X” symbol.

### 4.2 Schizophrenia RNA-seq dataset

The schizophrenia RNA-seq dataset (http://www.synapse.org/CMC) was obtained from the CommonMind Consortium [11] using post-mortem human dorsolateral prefrontal cortex tissues from 258 schizophrenia patients and 279 controls. Here we restricted our analysis to patients with age below 50 and post-mortem interval (PMI; time relapsed from the person has died to the tissues are frozen) less than 30 hours, producing 150 subjects where 100 are controls and 50 suffer from schizophrenia. Five clinical variables were available: age of death, gender, PMI, pH level and ethnicity (Caucasian or African American). At first, we ran a differential expression analysis on each covariate and found a higher percentage of DE genes affected by age of death, ethnicity, PMI and pH. However, since pH had some missing values, we only considered the other three clinical variables in the sNBLDA_GLM.sC_ model. We performed routine data preprocessing and filtering to keep genes with at least 70% of the samples having gene expression counts greater than 0 and mean count across the samples greater than 10, producing a count data matrix with 16989 genes for machine learning. Similar to simulation and previous application, 10-fold cross-validation was performed to evaluate sPLDA, NBLDA_PE_, sNBLDA_GLM_ and sNBLDA_GLM.sC_. We further performed DE analysis to narrow down to 50-1000 genes with a step of 50 in each training set before adopting the four machine learning methods. Even though three of the four methods have embedded feature selection capacity, the feature selection is usually difficult for ultra-high dimensionality (e.g. 16,989 gene features in our case). We performed a pre-screening by differential expression analysis to reduce dimensionality to 50-1000. This procedure is similar to the sure independence screening idea in [10] and can usually improve prediction performance.

Figure 4 shows the 10-fold cross validation accuracy of the four methods for different gene size after DE analysis pre-screening. For the three methods with embedded feature selection (sPLDA, sNBLDA_GLM_ and sNBLDA_GLM.sC_), varied tuning parameters for feature selection were applied and the best prediction accuracy was reported in Figure 4. The result clearly demonstrates better prediction performance of sNBLDA_GLM.sC_, especially when the pre-screening by DE analysis reduced the input gene size to 50-1000. However, when large number of genes were input to the sNBLDA_GLM.sC_ algorithm (e.g. more than 2000 genes after pre-screening), its performance dropped to close to sNBLDA_GLM_ and the advantage of covariate adjustment was diminished. Nevertheless, our proposed GLM approach generally outperformed sPLDA and NBLDA_PE_. As a result, we recommend pre-screening down to 50-1000 genes for ultra-high dimensional data, such as regular RNA-seq datasets, before applying sNBLDA_GLM.sC_. For completeness, we also compared sNBLDA_GLM.sC_ with sNBLDA_GLM.C_ (adjustment with all three covariates without sparsity of covariates) in Supplement Figure S3. The result shows inferior performance of sNBLDA_GLM.C_, indicating necessity of covariate regularization. We also performed SVM, RF and CART (Supplement Figure S5) and the prediction accuracies were generally much lower than sNBLDA_GLM.sC_ and sPLDA.

**Figure 4.**
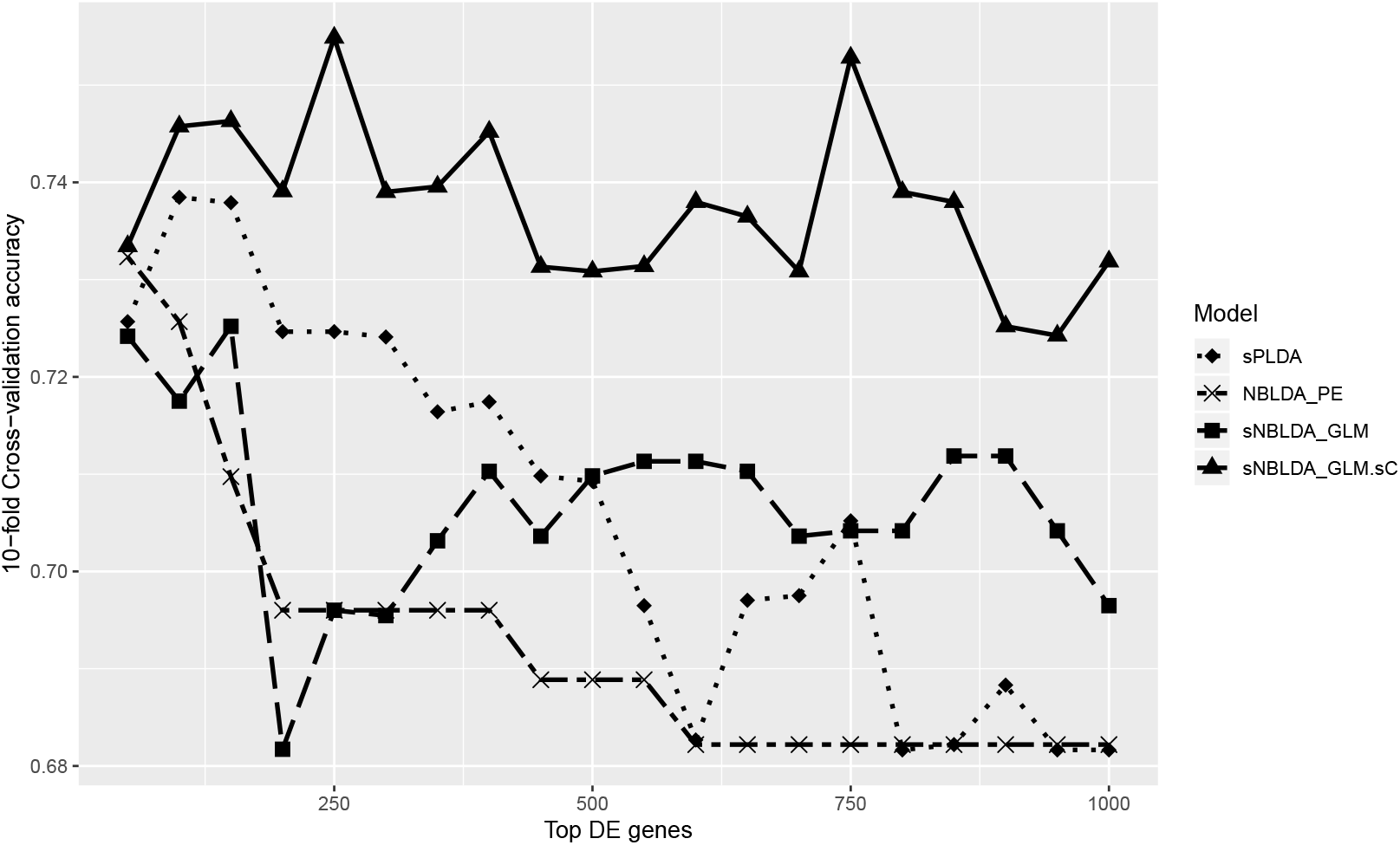
Prediction accuracy (y-axis) of sPLDA, sNBLDA_PE_, sNBLDA_GLM_ and NBLDA_GLM.sC_ with varying input gene number after DE analysis pre-screening (x-axis) in the schizophrenia post-mortem brain RNA-seq data.

## 5 Conclusion and Discussion

In this paper, we propose a sparse negative binomial classifier based on a GLM framework with and without covariate adjustment. The method incorporates three key elements in RNA-seq machine learning modeling: adequate modeling for count data, feature selection and adjustment of covariate effects. Existing methods such as sPLDA does not consider overdispersion properly, NBLDA_PE_ does not embed regularization for feature selection and both methods cannot adjust for covariate effect in gene expression. Our new approach assumes a negative binomial model to allow overdispersion, adopts GLM to allow covariate adjustment and facilitates double regularization for feature selection and covariate selection. Extensive simulations and two real applications showed superior performance of sNBLDA_GLM.sC_ in terms of prediction accuracy and feature selection. Particularly, our proposed method achieved higher prediction accuracy with smaller number of selected genes or miRNAs in the two real applications.

One major limitation of the four count data methods compared in this paper is that the methods are based on gene independent assumption. Due to the complex form of multivariate negative binomial model and the potentially heavy computational cost, it is not addressed in this paper but will be a future direction.

