## Supplementary_materials for "A sparse negative binomial classifier with covariate adjustment for RNA-seq data"

#### 1 Appendix A: Evaluation of Estimation of parameters under Simulation setting 2

The methods proposed in this paper follows a different estimation procedures compared to the the other two competing methods discussed in the this paper. It is important to note that the the estimation of the classification center mean for both sPLDA and NBLDA$_{\mathrm{PE}}$ are same since NBLDA$_{\mathrm{PE}}$ borrows the estimating strategy from the sPLDA. In particular, in this section we are interested to see how the precision of estimation of $\beta_{jk}$ is affected by the different level of strength of covariates on the the gene expression level. Under the formulation of sPLDA and our method, $\beta_{jk}$ in sNBLDA$_{\mathrm{GLM}}$ and and sNBLDA$_{\mathrm{GLM.sC}}$ is equivalent to $\log((g_j d_{jk})/n)$. We use the Simulation scheme 2 discussed in Section 3.1 with $\rho = 1$ , $\nu \in \{1, 10\}$ and $\eta \in \{0.05, 0.25, 0.45, 0.65, 0.85, 1.05, 1.25\}$. The result is summarized in Figure 1. As we can see with increasing covariate effect, the MSE increases sharply with both sNBLDA$_{\mathrm{GLM}}$ and sPLDA while it remains relatively constant in the case of sNBLDA$_{\mathrm{GLM.sC}}$. Under both scenario, we see that sNBLDA$_{\mathrm{GLM}}$ gives a more robust estimate compared to sPLDA.

#### 2 Appendix B: Advantage covariate selection in sNBLDA$_{\mathrm{GLM.sC}}$

When there is a strong signal in gene expression level, covariate selection is not quite important to in contrast to when the signal is very weak. Here, we run a simulation scheme following the steps in Simulation 2 scheme in Section 3.1 of the main paper. Here, we fix the $\rho = 0.125$ meaning that only 12.5% of the genes are actually affected by the covariate expression level. We vary the gene strength $\zeta \in \{0.15, 0.20, 0.35, 0.50\}$ on a range of the parameter $\nu \in \{0.5, 1\}$. We compare the performance of sNBLDA$_{\mathrm{GLM.C}}$ to sNBLDA$_{\mathrm{GLM.sC}}$ over different level of $\eta \in \{0.05, 0.1, 0.5\}$ and compare performance in terms of both testing accuracy and feature selection. The result is summarized in Figure 2. Here we see that the testing accuracy for sNBLDA$_{\mathrm{GLM.sC}}$ is higher compared to sNBLDA$_{\mathrm{GLMC}}$ for higher level of dispersion ($\nu = 0.5$). This suggests that with weaker signal, that is higher level of dispersion, covariate selection leads to higher level of testing accuracy. However, in terms feature selection, the performance of is comparable even though covariate selection can lead a small increase in the feature selection performance.

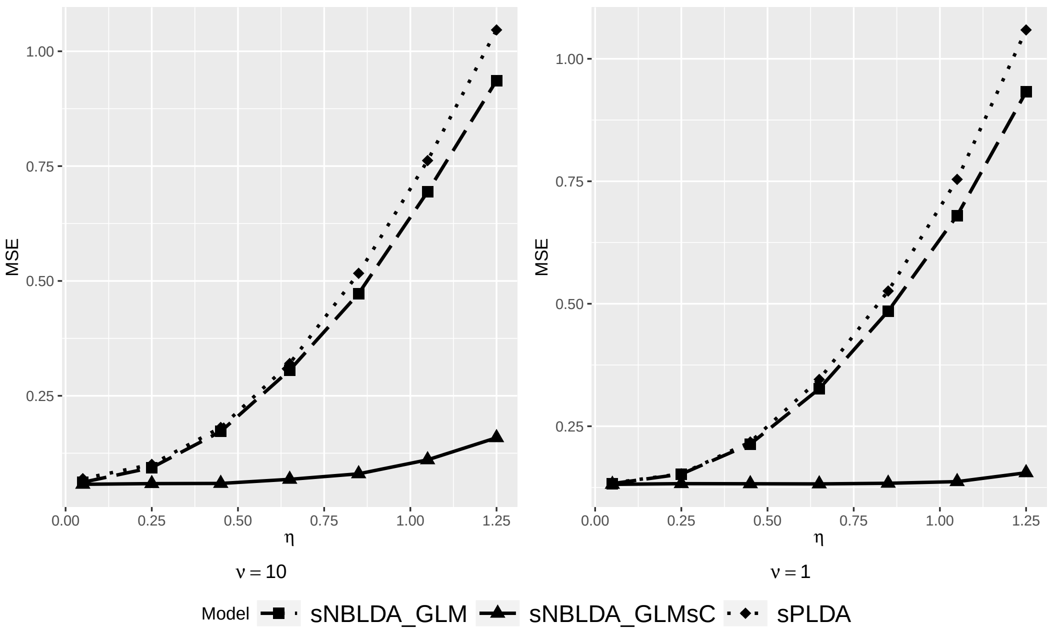


**Figure 1.** Evaluation of estimation of parameters with varying covariate effect and dispersion

In addition, we compare $\text{sNBLDA}_{\text{GLMC}}$ against $\text{sNBLDA}_{\text{GLM.sC}}$ in the schizophrenia dataset and the result is summarized in Figure 3. This clearly shows the advantage of using $\text{sNBLDA}_{\text{GLM.sC}}$ as compared to $\text{sNBLDA}_{\text{GLMC}}$ in terms of cross-validation accuracy. At all selection of top DE genes, $\text{sNBLDA}_{\text{GLM.sC}}$ outperforms $\text{sNBLDA}_{\text{GLMC}}$.

### 3 Appendix C: Comparison with some popular classifier used in case of continuous data

One common way to handle classification problem in RNA-seq is to convert the count data into continuous data and then apply methods developed for continuous data classification. Here, we compare the three methods discussed in the paper and our proposed model to the some of the more popular methods proposed in the literature used in classification problem. The methods considered here are random forest , support vector machine and carting denoted by RF, SVM and CART respectively. One popular method of transformaion in RNA-seq data is variable stabilizing transformation (VST). We compare the testing accuracy between these methods in Figure 4 using the Simulaton scheme 1 proposed in Section 3.1 of the paper. We find that the count based model outperforms all the the methods used in continuous variable case. Among, random forest, support vector machine and carting, random forest performs the best.

We also compared RF, SVM, CART with the methods discussed in presence of covariates affecting the gene expression level. The result is summarized in Figure 5. Here, we see that in presence of high dispersion, VST transformation ends up loosing information. As a result all the continuous data classification methods performs quite poor compared to the models designed specifically for the count data structure of the RNA-seq. However, in presence of low overdispersion ($\nu = 10$), we see the performance of random forest and support vector machine are comparable to $\text{NBLDA}_{\text{GLM}}$,

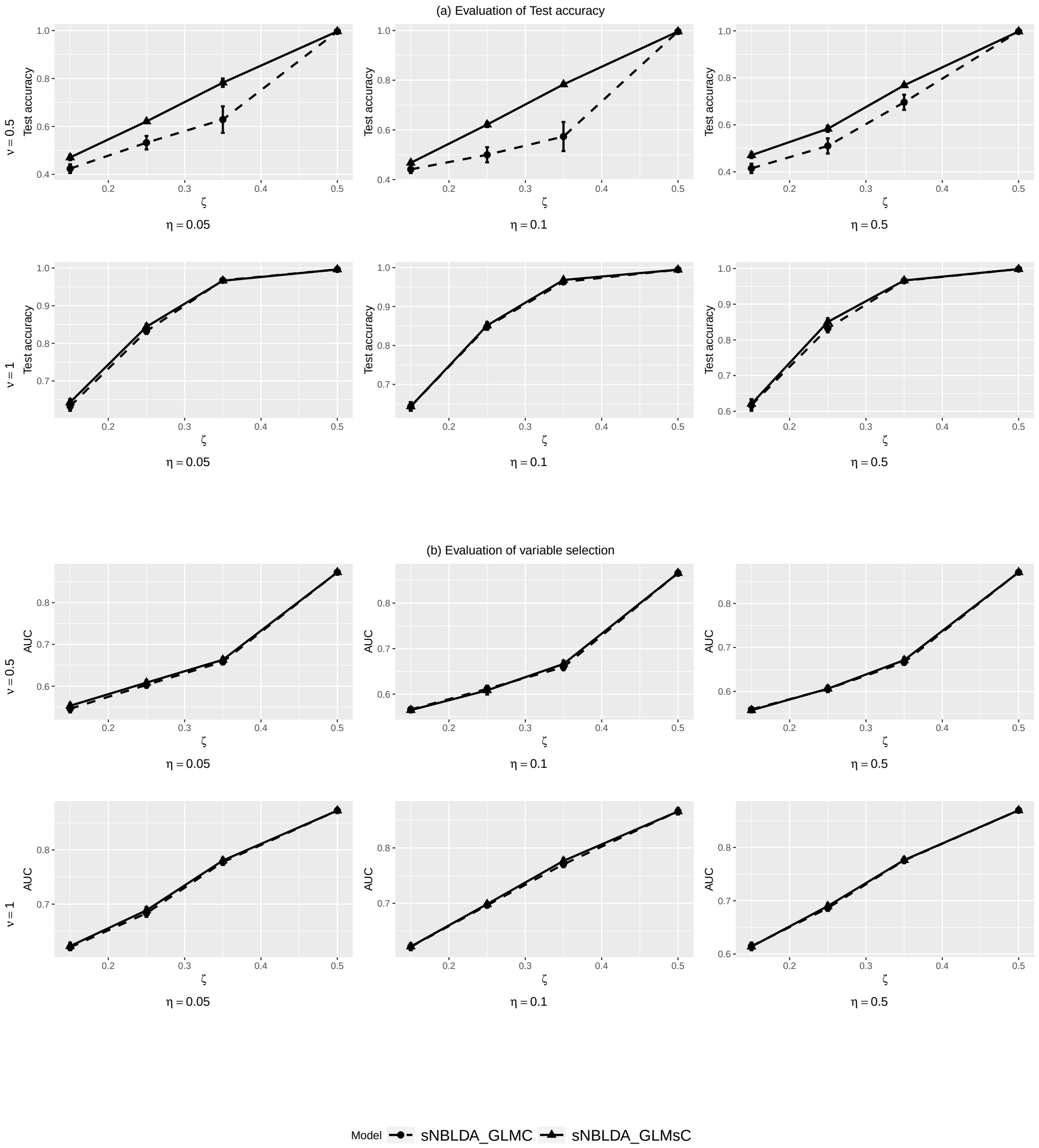


**Figure 2.** Comparison between covariate selection vs no covariate selection

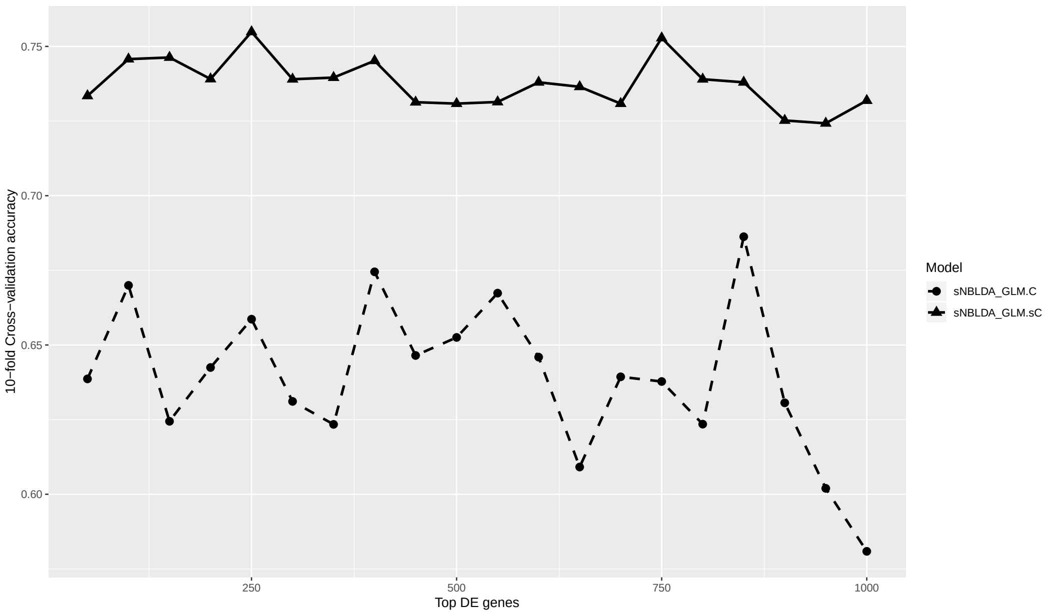


**Figure 3.** Comparison between covariate selection vs no covariate selection in Schizophrenia data

$\text{NBLDA}_{\text{GLM.sC}}$, sPLDA and $\text{NBLDA}_{\text{PE}}$ in presence of weak covariate effect. However, interestingly the performance of the continuous classification methods are found to have higher accuracy compared to $\text{NBLDA}_{\text{PE}}$ and sPLDA in presence of strong effect of the covariates on the gene expression level and is comparable to $\text{NBLDA}_{\text{GLM}}$ and $\text{NBLDA}_{\text{GLM.sC}}$. Perhaps this could be explained from the results summarized in Figure 1 where we see the estimates of parameters for sPLDA and $\text{NBLDA}_{\text{PE}}$ to be highly impacted by the covariate effects.

Next we compared RF, SVM and CART to sPLDA, $\text{NBLDA}_{\text{PE}}$ and $\text{sNBLDA}_{\text{GLM.sC}}$ in the case of cervical data. The result is summarized in Figure 6. The figure shows, the classification methods developed for the count data all significantly outperforms the methods applied on the VST transformation of the count data.

Similarly, 10-fold cross-validation accuracy is compared across the methods for Schizophrenia data and is summarized in Figure 7. Here, we see that the maximum accuracy overall is achieved by $\text{sNBLDA}_{\text{GLM.sC}}$. SVM classifier seems to be the second based classifier performong better than $\text{NBLDA}_{\text{GLM}}$, sPLDA and $\text{NBLDA}_{\text{PE}}$. One reasonable explanation is in real data, the data may not strictly come from negative binomial distribution.

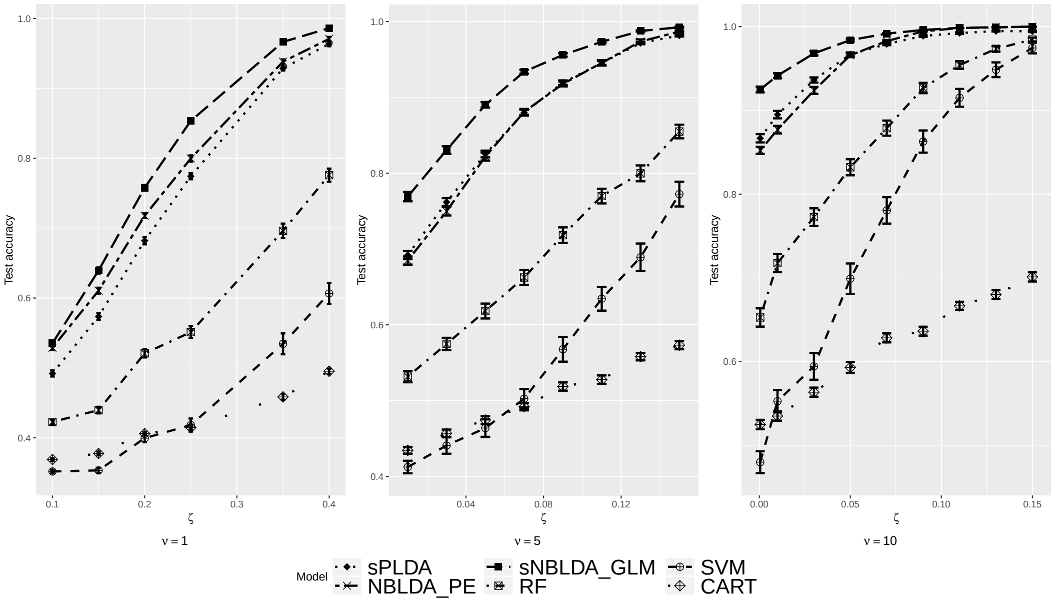


**Figure 4.** Evaluation of Testing accuracy in Simulation 1 scheme

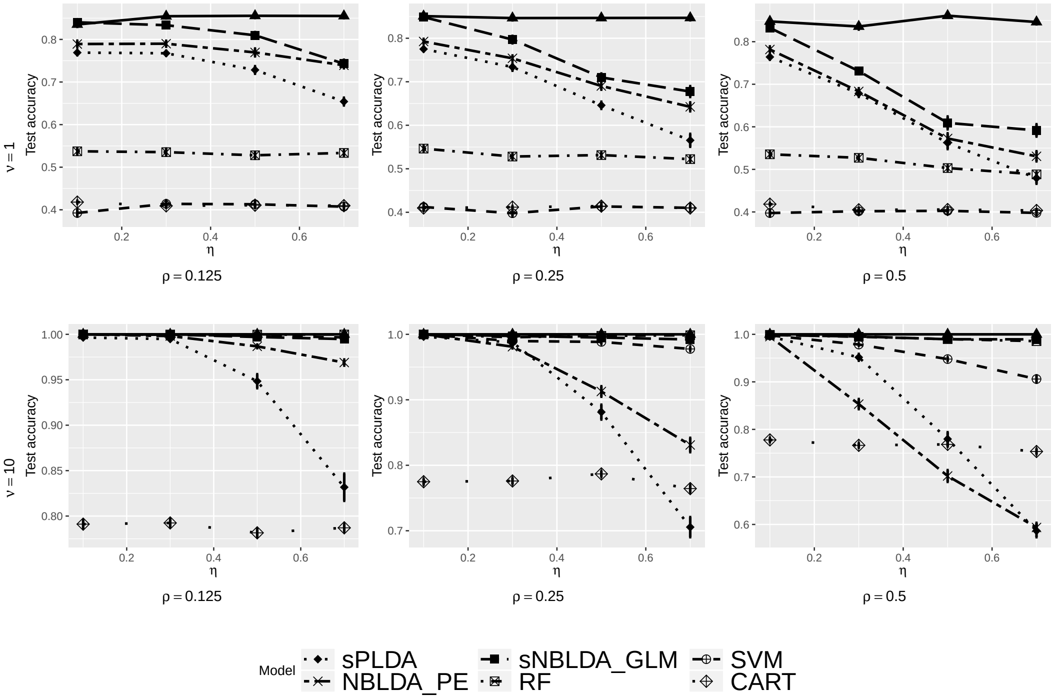


**Figure 5.** Evaluation of Testing accuracy under Simulation 2 scheme

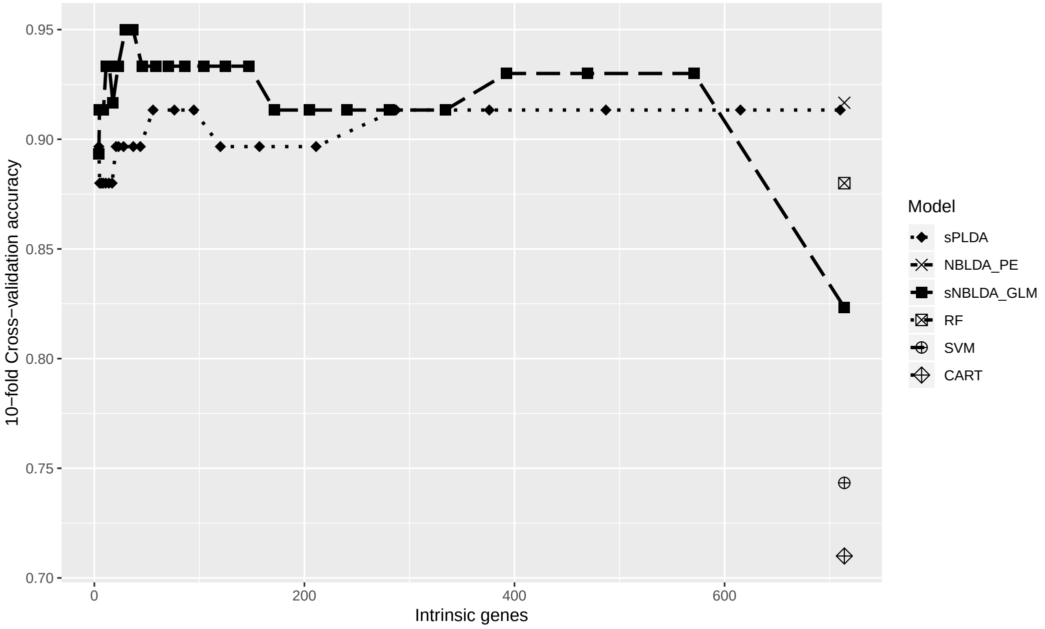


**Figure 6.** 10-fold cross-validation accuracy by Intrinsic genes

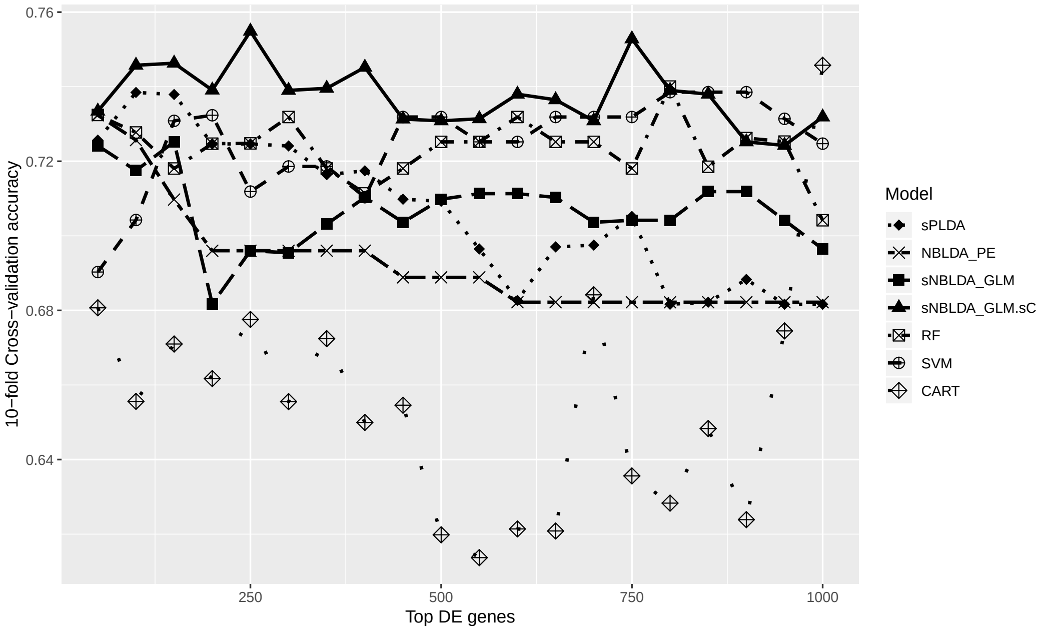


**Figure 7.** 10-fold Cross-validation by top DE genes in Schizophrenia data
